# Distinct representations of finger movement and force in human motor and premotor cortices

**DOI:** 10.1101/2020.02.18.952945

**Authors:** Robert D. Flint, Matthew C. Tate, Kejun Li, Jessica W. Templer, Joshua M. Rosenow, Chethan Pandarinath, Marc W. Slutzky

**Author notes:** Corresponding author: Robert D Flint, Research Assistant Professor, Neurology Feinberg School of Medicine Northwestern University.

## Abstract

The ability to grasp and manipulate objects requires controlling both finger movement kinematics and isometric force. Previous work suggests that these behavioral modes are controlled separately, but it is unknown whether the cerebral cortex represents them differently. Here, we investigated this question by recording high-density electrocorticography from the motor and premotor cortices of seven human subjects performing a sequential movement-force motor task. We decoded finger movement (0.7±0.3 fractional variance account for; FVAF) and force (0.7±0.2 FVAF) with high accuracy, yet found different spatial representations. We also found clear distinctions in electrocorticographic activity by using deep learning methods to uncover state-space representations, and by developing a new metric, the neural vector angle. Thus, state-space techniques can help to investigate broad cortical networks. Finally, we were able to classify the behavioral mode from neural signals with high accuracy (90±6%). Thus, finger movement and force have distinct representations in motor/premotor cortices. This will inform our understanding of the neural control of movement as well as the design of grasp brain-machine interfaces.

## Introduction

The human ability to grasp and manipulate objects is central to our evolutionary success as tool users. The loss of this ability has a profound negative impact on overall quality of life. We rely in particular upon our ability to precisely regulate movement and force, to close our fingers around an object, then exert isometric force sufficient to prevent slippage without crushing it. However, the neural origin of this process is not yet clear. In the current study, we sought to identify how (or whether) movement and force are encoded differently at the cortical level.

There is longstanding evidence for cortical representations of both movement (Moran and Schwartz, 1999) and force (Evarts, 1968). Further, there is indirect evidence that distinct neural control states are used for kinematics (movement) and kinetics (force). For example, motor learning of kinematics and kinetics in reaching occur independently of each other (Flanagan et al., 1999). Kinematic and kinetic control can be disrupted independently (Chib et al., 2009), and their errors can be separated during adaptation (Danion et al., 2013). Perhaps most relevant, Venkadesan and Valero-Cuevas (2008), found that electromyogram (EMG) activity patterns transitioned between separate, incompatible states during a one-finger, sequential movement-force task. Importantly, these transitions occurred prior to the fingertip’s contact with a surface, implying that changing neural states may “prepare” finger muscle activations for their upcoming role in regulating force. Here, we hypothesized that the transition between movement and force is encoded in motor and premotor cortical networks.

The specifics of cortical movement and force encoding are also relevant to brain-machine interface (BMI) design. Restoration of hand grasp functionality is a high priority for individuals with paralysis (Blabe et al., 2015). Currently, BMIs using motor cortical signals control robotic or prosthetic hands (Hochberg et al., 2012; Yanagisawa et al., 2012; Wodlinger et al., 2014; Hotson et al., 2016), or functional electrical stimulation of paralyzed limbs (Pfurtscheller et al., 2003; Bouton et al., 2016; Ajiboye et al., 2017). However, most BMIs that have decoded grasp intent have focused on decoding kinematics of grasp aperture. One exception improved BMI-prosthetic hand control by scaling the neuronal firing rates (Downey et al., 2017), but did not examine the movement-force transition. Here, we hypothesized that force and kinematics of the hand are governed by separate neural states in cortex.

In the current study, we used a sequential movement-force task to investigate changes in human cortical activity during transitions in behavioral mode: from pre-movement (preparation) to movement to force. We recorded subdural surface potentials (electrocorticography; ECoG), finger kinematics, and applied force. We used ECoG spectral modulations to measure changes in the spatial patterns of movement- and force-based decoding, and to classify the behavioral mode of the subject. We found clear evidence of distinct movement and force encoding.

Recent work has characterized changes in cortical network activity during kinematic tasks as the temporal evolution of a dynamical system (Churchland et al., 2012; Pandarinath et al., 2018). Here, we examined whether neural state space changes accompanied behavioral mode transitions (from pre-movement to movement to force). We used latent factor analysis via dynamical systems (LFADS), a deep-learning method that uses sequential autoencoders to uncover trajectories in a low-dimensional neural state space from high-dimensional neural data (Pandarinath et al., 2018). We also calculated changes in a neural vector angle (NVA), obtained by treating the spectral features as elements of a high-dimensional neural vector. Both approaches showed that activity across a broad area of motor and premotor cortices exhibited tightly clustered trajectories through neural state space that were time-locked to the behavior. The NVA enabled us to average responses across subjects and create a generalized temporal profile of neural state space activity during the movement and force modes of human grasp. Together, these analyses indicate that distinct cortical states correspond to the movement and force modes of grasp.

## Materials and Methods

### Subjects and recordings

Seven human subjects participated in the study (all male; ages 26-60, ordered chronologically). Six of the subjects required awake intraoperative mapping prior to resection of low-grade gliomas. Their tumors were located remotely to the cortical areas related to hand grasp, and no upper extremity sensorimotor deficits were observed in neurological testing. Subject S6 underwent extraoperative intracranial monitoring prior to resection surgery for treatment of medication-refractory epilepsy. All experiments were performed under protocols approved by the institutional review board of Northwestern University. All subjects gave written informed consent before participating in the study. Subjects were recruited for the study if the site of their craniotomy, or their monitoring array was expected to include coverage of primary motor cortex.

In all subjects except S6, we used 64 electrode (8×8) high-density ECoG arrays, with 1.5-mm exposed recording site diameter and 4-mm inter-electrode spacing (Integra, Inc.). Arrays were placed over hand motor areas, which we defined by: 1) anatomical landmarks, e.g., ‘hand knob’ in primary motor cortex; 2) pre-operative fMRI or transcranial magnetic stimulation to identify functional motor areas; and 3) direct electrocortical stimulation mapping. Intraoperative recordings took place after direct stimulation mapping. Intraoperative MRI navigation was performed with Curve (BrainLab, Inc., Munich, Germany). The recording arrays covered primary motor cortex, premotor cortex, and usually part of primary somatosensory cortex as well (Figure 1A). In S6, electrode placement was determined by clinical need. For this subject, we used a 32-electrode (8×4) array with the same electrode size and spacing as our 64-electrode arrays.

**Figure 1.**
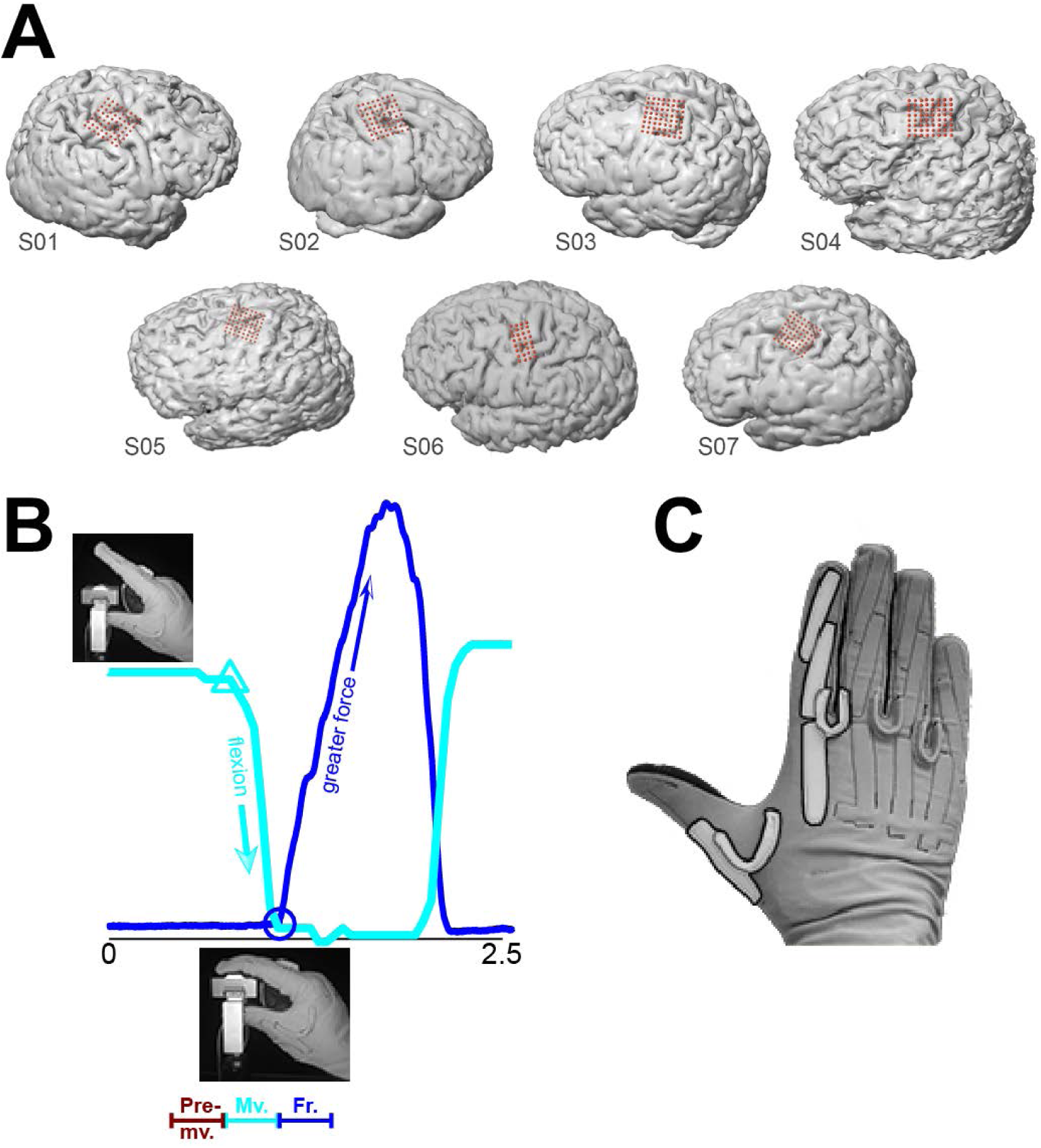
ECoG array placement, experimental task, and behavioral data. (**A**) In S1 through S5 and S7, we targeted the primary motor and premotor cortices. Array placement for S6 was determined by clinical need. For S1 and S2 we recorded ECoG from the right hemisphere; the other subjects’ ECoG were recorded from the left hemisphere. (**B**) One trial (approximately 2.5s) of the kinematic-kinetic task. At the beginning of the trial, the subjects held their index finger in a neutral position (upper left photograph) until visually cued on a screen. Cyan trace: finger kinematics (amount of flexion; arbitrary units) during the trial. Cyan triangle: time of flexion movement onset. Upon contact with the force sensor (lower inset photograph), the subjects exerted isometric force until matching a force target on the screen with a cursor (not shown). Blue trace: recorded force. Blue circle: time of force onset. At bottom is a schematic representation of behavioral mode segmentation: pre-movement (from target presentation until the start of flexion), movement (start of flexion until start of force), and force (from force onset lasting 500ms). (**C**) We measured index finger flexion using a CyberGlove; movement onset was identified using the first principal component calculated on the data from the highlighted sensors.

We sampled ECoG at 2 kHz using a Neuroport Neural Signal Processor (Blackrock Microsystems, Inc.). Signals were bandpass filtered between 0.3 Hz and 500 Hz prior to sampling. Finger kinematics were recorded using a 22-sensor CyberGlove (Immersion). We recorded force with a custom-built load cell sensor. Kinematic and kinetic data were both sampled at the same rate as ECoG.

### Experimental protocol

The subjects executed repeated trials of a one-finger task that required isotonic movement and isometric force in sequence. At the beginning of each trial, the subjects were instructed to hold their index finger in a neutral posture. After a cue, they executed a self-paced flexion movement (Figure 1B), which brought the palmar surface of the index finger into contact with the force sensor. Upon contact, subjects were instructed to apply force to the sensor, thereby controlling a cursor on a monitor. Their task was to match the cursor’s vertical position to that of a force target presented on the monitor. Target force levels varied randomly from trial to trial (random-target pursuit task, as in Flint et al., 2014). Following a successful match (or a timeout of 2s), the trial was complete, and the subject extended their finger back to the baseline (neutral) position. The next trial began after a delay of 1s. Target presentation and cursor feedback were controlled by the open-source BCI2000 software (Schalk et al., 2004). The time resolution for both kinematic data acquisition and force cursor control was 50ms.

Our task was designed to elicit movement in one finger, while keeping the other fingers motionless in a flexed position. Therefore, we analyzed only the data from the CyberGlove sensors that were relevant to the motion of the index finger (Figure 1C, highlighted). Dominant kinematic features were extracted via principal component analysis (PCA), similar to (Flint et al., 2017). We performed PCA only on data from the highlighted sensors in Figure 1C, retaining the 1^st^ component to identify movement onset.

### Feature extraction

For all analyses, we extracted spectral features from each ECoG electrode. Here, each feature was the mean spectral power in a frequency band of interest. These methods followed closely from our published studies of decoding isometric force (Flint et al., 2014) and movement kinematics (Flint et al., 2017) from ECoG. We calculated the log-normalized spectral power in each ECoG electrode using short-time Fourier transforms (window width of 512 ms). We averaged the spectral power in 25-ms time bins. We identified the feature boundaries (frequency bands of interest) by computing the event-related spectral perturbation (ERSP) for each electrode around the time of force onset. We then averaged the ERSPs for all electrodes in our dataset, and identified the frequency bands of interest: broadband low frequency (8-55Hz) and broadband high frequency (70-150Hz). Subsequent analyses were performed on the feature matrix for each subject. Each feature matrix was size NxM, where N is the number of time bins in the record, and M is 2*(number of electrodes)*10, where 10 was the number of time bins into the past (causal bins only).

### Population decoding of continuous movement and force

We decoded continuous movement kinematics and continuous isometric force, using all (non-noisy) electrodes from PM and M1 in each subject. For continuous decoding, the feature matrix served as input to a Wiener cascade decoder (Hunter and Korenberg, 1986). In the Wiener cascade, the output of a linear Wiener filter is convolved with a static nonlinearity (here, a 3^rd^-order polynomial). We employed ridge regression to reduce the likelihood of overfitting due to the large feature space, as in (Suminski et al., 2010). We evaluated decoding accuracy using the fraction of variance accounted for (FVAF). We employed 11-fold cross-validation, using 9 folds for training, 1 fold for parameter validation (e.g., optimizing the free parameter in the ridge regression Fagg et al., 2009), and 1 fold for testing. We report the median ± interquartile range (IQR) of FVAF across test folds.

### Spatial mapping of decoding performance

We quantified the difference in the spatial representations of movement and force using two measures: (1) change in location of the peak single-electrode decoding performance, and (2) change in the overall spatial distribution of single-electrode decoding performance. For both analyses, we decoded continuous movement for each individual ECoG electrode using Wiener cascade decoders, as in the previous section. These single-electrode decoding results were evaluated using the cross-validated FVAF, as above. The spatial distribution of single-electrode movement decoding performance formed a “map” for the array. In a similar manner, we constructed a “map” of force decoding performance. We then analyzed these maps to reveal differences between movement and force, in terms of spatial representation on the cortical surface.

We compared the location of the overall peak of each decoding map for movement to that of force within each cross-validation fold. We report the absolute displacement between the peak performance location from force decoding vs. that from movement decoding. Peak performance displacement quantifies the shift in location between movement and force in units of distance (e.g., in millimeters).

In addition, we compared the overall decoding map patterns. The map for a single fold can be treated as an image, with FVAF values corresponding to pixel intensities. We measured similarity among maps using a differencing metric common to image processing (Euclidean distance). We calculated the distance (D) between pairs of maps for individual folds. For example, a value of D_intra,3-4(force)_=0, where D is the difference metric, would indicate that the force decoding maps in folds 3 and 4 were identical. We compared the inter-map distances across behavioral modes (movement vs. force, D_inter_) to find the average decoding map difference between movement and force encoding on the cortex. We compared these to within-modality distances (D_intra(force)_,D_intra(mvt)_), which vary only due to time. That is, D_intra_ measured map differences within a behavioral mode, which can be attributed to variance in task performance across trials. Thus, D_intra_ values served as controls for D_inter_, which measured the map differences attributable to control mode (movement or force). When calculating these distance metrics between performance maps, we scaled by the maximum possible distance between the maps, so that both D_inter_ and D_intra_ ranged from 0 to 1.

### Latent factor analysis via dynamical systems

We applied a deep learning algorithm, latent factor analysis via dynamical systems (LFADS), to denoise ECoG features (Sussillo et al., 2016; Pandarinath et al., 2018). LFADS attempts to denoise neural activity based on the assumption that the observed patterns of neural modulation can be described as noisy observations of an underlying low-dimensional dynamical system. LFADS aims to extract a set of low-dimensional latent factors that describe neural population activity on a single-trial basis. When previously applied to spiking activity from populations of neurons, LFADS modeled observed spikes for each neuron as samples from an inhomogeneous Poisson process (called the firing rate), and attempted to infer this underlying firing rate for each neuron. In this study, since the ECoG features are continuous rather than discrete variables, the underlying distribution was taken to be Gaussian instead of Poisson.

Specifically, we first pre-processed the data by z-scoring each spectral feature. We then modeled the data following the equations in Sussillo et al. (2016), with the key modifications that:

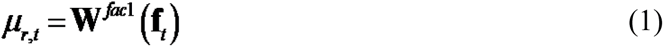

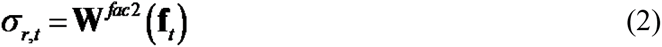

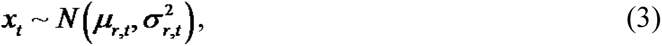

where **x***_t_* represents the vector of z-scored spectral features at each timestep, and **f***_t_* represents the latent factors output by the LFADS recurrent neural network. For a given spectral feature *r*, *μ_r,t_* and *σ_r,t_* represent the inferred time-varying mean and variance, respectively, for the z-scored spectral feature at each time step. *W^fac^*^1^ and *W^fac^*^2^ are matrices that map the latent factors onto *μ_r,t_* and *σ_r,t_*, respectively. These matrices have fixed weights across all time points. For each subject, the number of latent factors allowed was approximately half the total number of ECoG channels used. After applying LFADS, we used principal component analysis to produce low-dimensional visualizations of the denoised ECoG features.

### Neural vector angle

To compactly represent the overall response of a subject’s feature set, we computed neural vector angles (NVAs) for each trial. This quantity is similar to the “muscle coordination pattern” angle of Venkadesan and Valero-Cuevas (2008). We selected features to include in the NVA calculations using the following method: first, we averaged the spectral intensity across trials, aligned to force onset. We then used unsupervised k-means clustering (3 clusters) to partition the trial-averaged spectral power from the complete set of features. For a subject with 64 non-noisy electrodes, this would mean that 128 features (64 low-frequency features, 64 high-frequency features) served as the inputs to the clustering algorithm. Of the three output clusters, we selected the two that were well-modulated with movement and/or force: a cluster of low-frequency features and a cluster of high-frequency features. These groupings (low- and high-frequency features) emerged natively from the unsupervised procedure. Clustering was used only as a means of selecting ECoG features to include in NVA computations.

We calculated the NVA for the selected features in each cluster as follows: a cluster of features with n members can be represented at time t as **m**(t)=[f_1_,f_2_,…,f_n_], where f is the value of an individual feature. We smoothed **m**(t) over 5 time bins (total 125 ms), then calculated the neural vector angle

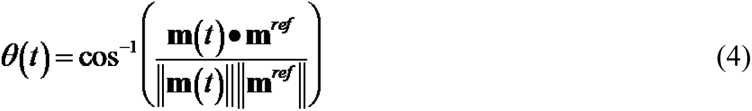

where **m**^ref^ is the average value of **m**(t) over the 250-ms period before the time of maximum force exertion in the trial. We computed the neural vector angle at each time bin over trials in each of the emergent clusters (low- and high-frequency modulating), for each subject. Since the neural vector angle transformed the data from feature values to a common coordinate system (angle between vectors, in degrees), it enabled us to average this quantity across subjects. To quantify differences in NVA values due to behavioral mode, we used the Kruskal-Wallis test of unequal medians on NVAs during “pre-movement”, “movement”, and “force” modes (Figure 1B). See also the following section for details of the behavioral mode labelling procedure.

### Discrete classification of behavioral mode

Our classification of behavioral mode used the same feature matrix as continuous decoding. Data were labeled as follows: time points from the time of target presentation to the start of finger flexion were labeled as “pre-movement”; time points from the start of flexion to contact with the force sensor were labeled “movement”; time points beginning at contact with the force sensor, continuing for 0.5 s were labeled “force”. We limited the length of the force window to obtain more balanced class sizes. Data outside of the described time windows were discarded. The remaining data were classified using two methods: support vector machines and boosted aggregate (bagged) trees. The classification analyses used 5-fold cross validation. Within each test fold, we classified every 25-ms time bin. The reported accuracy measures are the median ± IQR of correctly classified time points across all test folds. Because the class sizes were not exactly equal, the chance level performance of the 3-class classifier was not necessarily 1/3. We calculated the true chance level performance by shuffling the class labels and then performing the analyses as above. We repeated this procedure 1000 times for each recording.

### Experimental design and statistical analysis

We conducted the experiments and analyzed the data using a within-subject design. We used non-parametric statistics to report continuous kinematics and continuous force decoding accuracy, as the decoding accuracy values (FVAF) were distributed non-normally across cross-validation folds. To compare maps of decoding performance, we conducted a one-tailed Wilcoxon signed-rank test, with Bonferroni correction for multiple comparisons. Differences in NVA during behavioral modes were tested using a Kruskal-Wallis test. For the discrete decoding of behavioral mode, we also used a Kruskal-Wallis test to identify statistical differences between ECoG feature-based decoding and LFADS-cleaned feature decoding.

## Results

We recorded ECoG from seven human subjects with brain tumors or epilepsy who required intraoperative or extraoperative mapping as part of their clinical treatment. In all subjects, ECoG coverage included at least part of primary motor and premotor cortices (Brodmann areas 4 and 6). In some cases, coverage also included prefrontal and/or postcentral cortices (Figure 1A). However, we restricted our analyses to electrodes covering primary motor and premotor cortices. The subjects performed a cued one-finger task requiring an isotonic flexion movement, followed by isometric flexion to specified force targets. Movement and isometric flexion were performed sequentially (Figure 1B). This task (adapted from Venkadesan and Valero-Cuevas, 2008) activates the same flexor muscles to achieve two different aspects of object grasp. We recorded the finger joint kinematics (based on the sensors highlighted in Figure 1C) as well as the force generated by isometric flexion.

### ECoG feature modulations were time-locked with movement and force

Following Collard et al. (2016), we used event-aligned plots to visualize event-related changes in ECoG spectral features, specifically to understand how tightly these features modulated with behavioral events. We examined modulation with respect to (1) the start of finger flexion movement and (2) the start of isometric force exertion. We constructed the intensity raster for each feature by windowing its data, then plotting as trial number vs. peri-event time. We sorted trials by the time between events.

We constructed raster plots for each feature in our dataset (2 features per non-noise electrode, 722 total features in the dataset). Overall, we found a diverse set of activity patterns during movement and force production. In the high-frequency range, spectral power increased around the start of isometric force, differentiating force production from movement (Figure 2A-C). Figure 2A shows an example of a high frequency feature that differentiated finger movement from both rest (Figure 2A, left of dashed line) and force (right of blue circles). Other high frequency feature modulations were time-locked only to force execution (Figure 2B,C), not to movement. By contrast, low frequency features (Figure 2D-F) showed mostly power decreases at movement onset. However, on occasion low-frequency power decrease was time-locked to the start of force, instead (Figure 2F). Note that Figures 2B and 2E show high- and low-frequency features from the same ECoG electrode, illustrating that two motor control behavioral modes can be encoded differently by high- and low-frequency information on the same electrode. We would encounter this trait again on a wider population level, during our neural vector angle analysis (later in Results).

**Figure 2.**
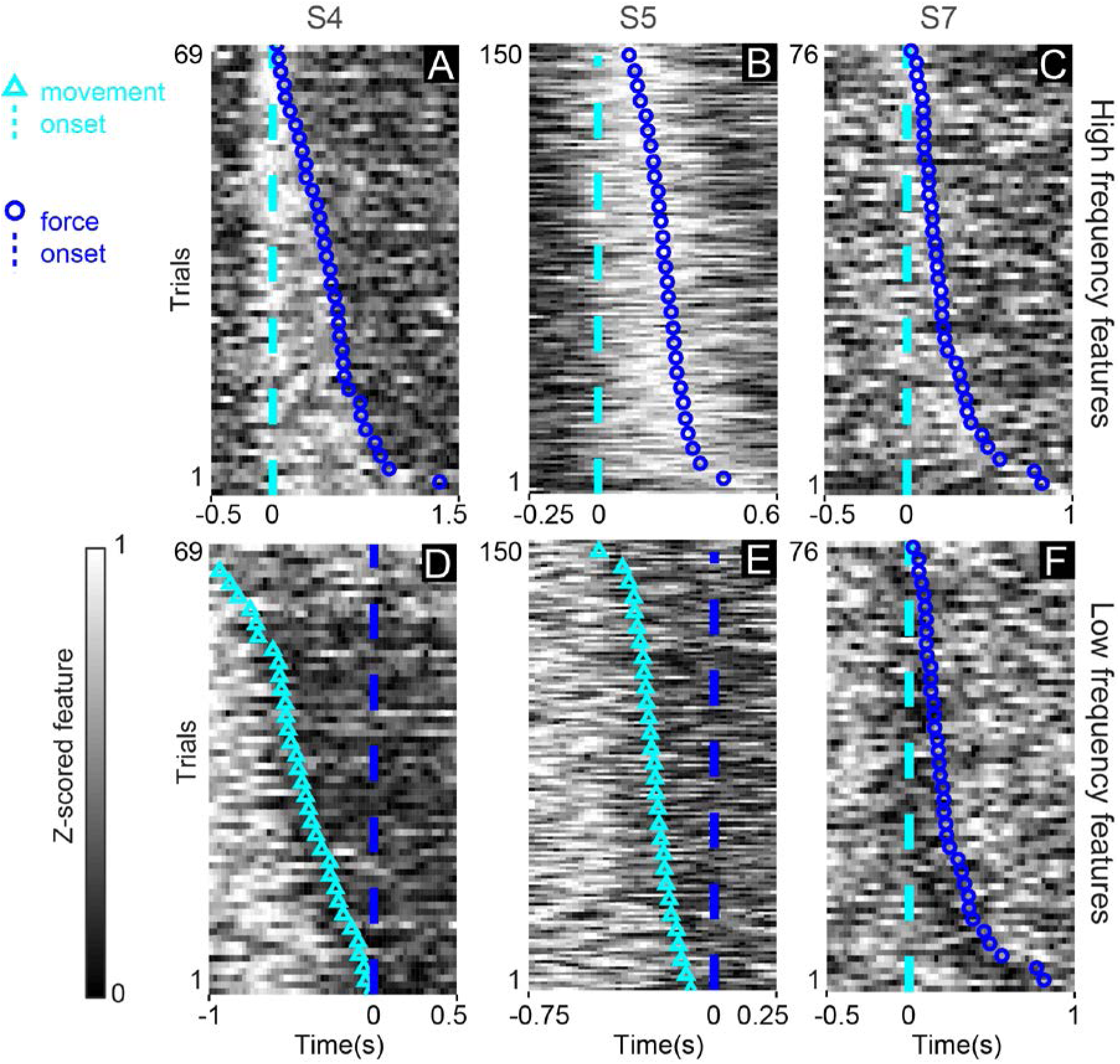
Spectral power modulation during the movement-force grasp task. Each panel shows data from a high- or low-frequency spectral feature taken from an individual ECoG electrode. The single-trial frequency band power (grayscale in each plot) was z-scored and aligned either to movement onset (cyan dashed lines, **A**-**C**,**F**) or to force onset (blue dashed lines, **D**-**E**). Blue circles show force onset times when trials were aligned to movement onset. Cyan triangles show movement onset times when trials were aligned to force onset. High frequency features (**A**-**C**) exhibited power increases, which could be time locked to both movement and force (**A**) or force only (**B**,**C**). Low frequency features (**D**-**F**) exhibited power decreases just preceding, and aligned to, the onset of movement (**D**,**E**), or aligned to the start of force (**F**).

### Continuous movement and force were decoded with high accuracy

Similar to our previous studies, we used a Wiener cascade decoder to build multi-input, single-output models for decoding behavior. We used one such model to decode the continuous time course of finger movement kinematics using both high and low spectral features from all (M1/PM) electrodes. A separate model was used to decode continuous isometric force from the same electrodes. The resulting decoding accuracy was high for both force and kinematics: the fraction of variance accounted for (FVAF) ranged from 0.4±0.1 (median±IQR) to 0.8±0.1. Across subjects, the overall median FVAF was 0.7±0.2 for force decoding, and 0.7±0.3 for movement decoding. Statistically, the null hypothesis that movement kinematics and force were decoded with equivalent accuracy could not be rejected (Kruskal-Wallis test, p=0.6); thus, our ability to distinguish between movement and force (reported in the following sections) was not due simply to decoding one quantity better than the other.

### Spatial mapping of decoding performance shows different cortical representations of movement and force

We next quantified the difference in the spatial representations of force and movement on the cortical surface, using two metrics: (1) change in location of the peak decoding performance electrode (Table 1), and (2) change in overall map pattern (Figure 3).

**Figure 3.**
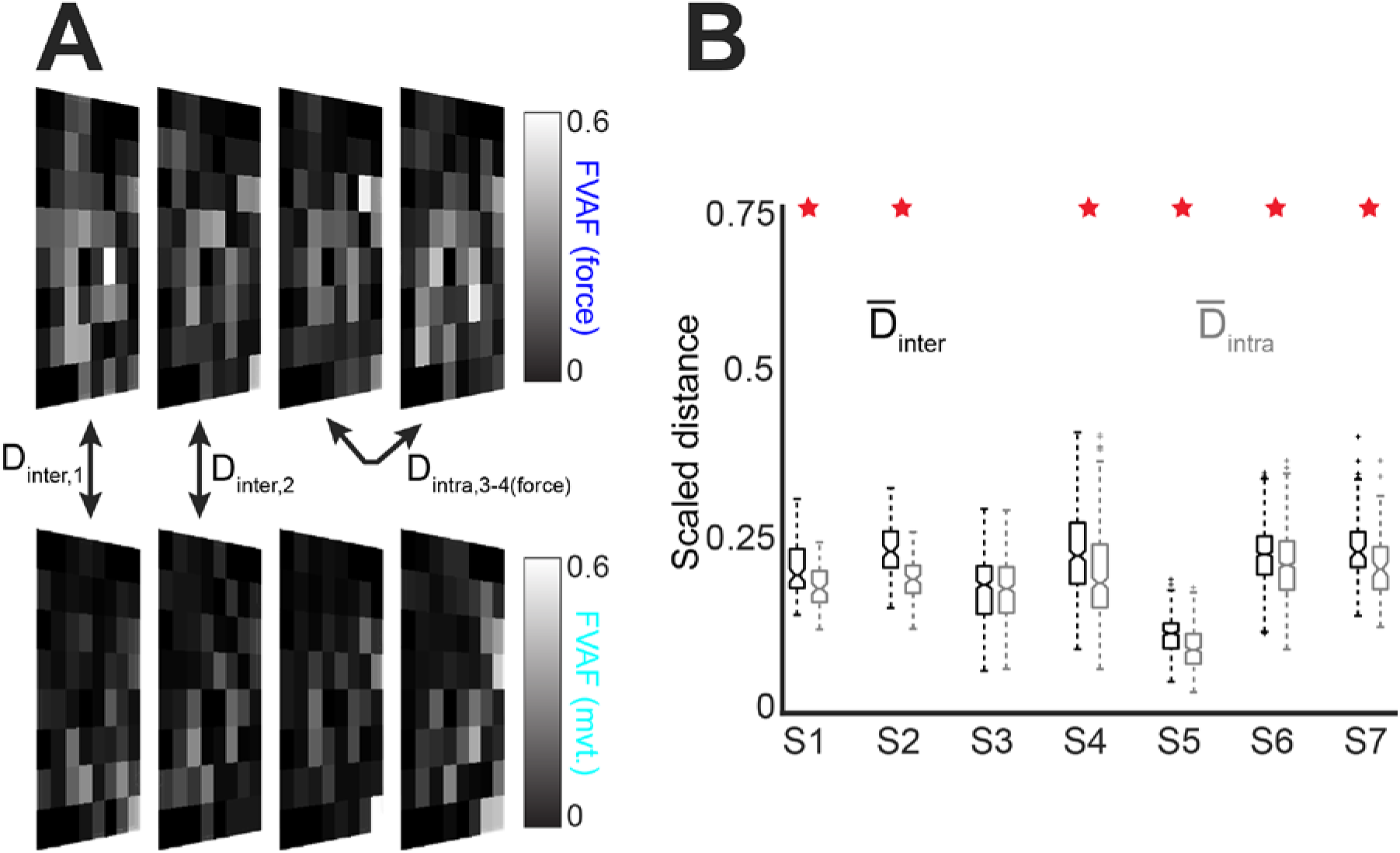
Decoding maps reveal changes in the cortical representations of movement and force. (**A**) Example decoding maps for 4 folds of data (the actual analysis utilized 10 folds per recording). Square recording arrays are shown in a rotated perspective for compact visualization. We compared single-electrode decoding maps for movement (top) and force (bottom) using a distance metric *D_inter_* for every possible combination of fold pairs. As a control, we calculated *D_intra_* between all possible fold pairs, for within-movement and within-force decoding. (**B**) Boxplot of distance measures for all subjects. The central horizontal line in each box shows the median, while the notches show 95% confidence intervals. Overall, the median *D_inter_* was significantly greater than the median *D_intra_* in 6 of 7 subjects (red stars).

**Table 1.** Displacement of peak location for movement decoding performance relative to force decoding performance in each subject.

| | mean | ± | S.D. |
| --- | --- | --- | --- |
| S1 | 16.1 | ± | 4.1 |
| S2 | 16.5 | ± | 8.8 |
| S3 | 3.2 | ± | 5.4 |
| S4 | 10.2 | ± | 8.4 |
| S5 | 4.2 | ± | 6.6 |
| S6 | 8.8 | ± | 5.4 |
| S7 | 10.7 | ± | 8.0 |

We previously showed that peak performance location differs for an isometric force performed with two different fingers (Flint et al., 2014). Here, we found that the peak performance location was different for movement and force decoding. The displacement (between movement and force) of the peak decoding performance ranged from 3.2±5.4 mm to 16.5±8.8 mm across subjects (mean ± SD over folds; Table 1). The mean (±SE) displacement of peak performance for all subjects was 9.9±2.0 mm.

To place these distances in context, a standard ECoG array for epilepsy use has an inter-electrode distance of 10 mm, highlighting the advantages of using high-density ECoG arrays (the electrode arrays used here had an inter-electrode distance of 4 mm). See also Wang et al. (2016).

In addition to changes in peak decoding location, there were differences between movement and force in the overall map patterns (Figure 3). The between-mode distance D_inter_, which measured differences between the movement-force maps (see Methods), was significantly greater than the within-mode distance D_intra_ in 6 of 7 subjects (p<3*10^-5^ except S3, where p=0.19; one-tailed Wilcoxon signed-rank test with Bonferroni correction for multiple comparisons; see Figure 3B).

This indicates that the spatial distribution of decoding as a whole changes between movement and force, and that this change is greater than what is expected from behavioral variation.

Taken together, these results indicate that the spatial representations of movement and force on the cortical surface are different. This difference was observable both in the location of peak decoding performance, as well as in the decoding map changes between behavioral modes.

### Differences in pre-movement, movement, and force behavioral modes are reflected in a dynamical systems model of cortical network activity

We next examined the activity of the recorded cortical network as a whole during the movement-force behavior. The preceding spectral/spatial analyses treated individual ECoG electrodes as independent sources of information. Here, we instead sought a low-dimensional representation to clarify and summarize the activity of the cortical network during the time course of the behavior. We used latent factor analysis via dynamical systems (Pandarinath et al., 2018) to generate low-dimensional representations of single-trial activity in the ECoG feature state space (see Methods). To visually summarize the factors, we compute principal components of the LFADS-denoised ECoG features (labeled LFADS-PCs). Figure 4 shows the underlying dynamics for S5 and S6 during trials of the kinematic-kinetic task, color-coded by three behavioral modes. At the start of the task (pre-movement), the high- and low-frequency latent factors tended to be distributed through a relatively broad region of the state space (ex. Figure 4A, red). Prior to the start of movement, the latent factors tended to converge onto a smaller region of state space, and their trajectories through the movement (cyan) and force (blue) periods of the task were more tightly grouped. Moreover, each time period of the task occupied a different part of state space (note the grouping of colors in Figure 4). To illustrate the impact of LFADS in revealing well-ordered, low dimensional state space representations, we also performed PCA directly on the ECoG features (PCA-only; Figure 4, inset boxes).

**Figure 4.**
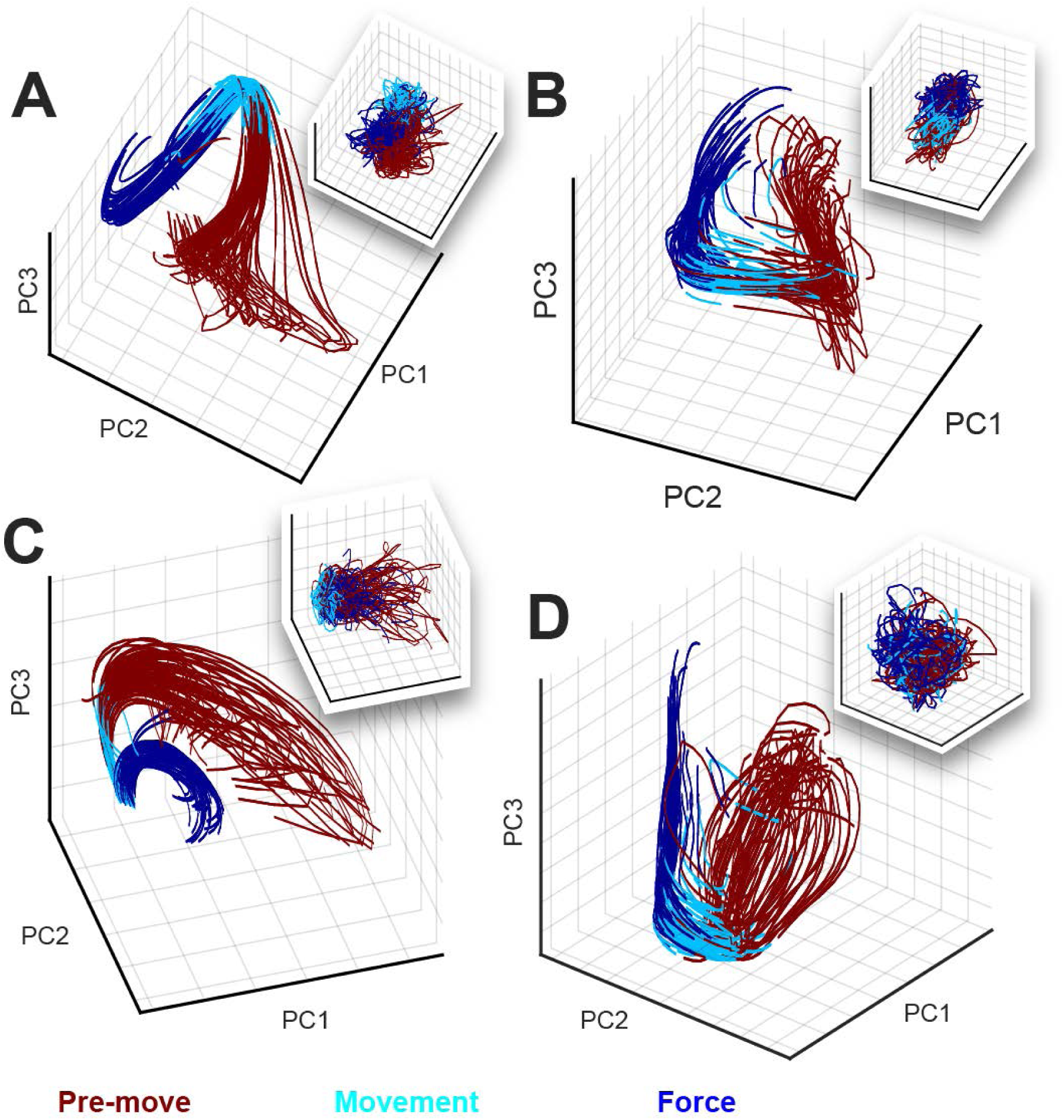
Modeling ECoG features as an underlying dynamical system using LFADS uncovers repeatable trajectories through a low-dimensional state space during the kinematic-kinetic task. Shown are LFADS-PCs (labeled as “PC” for simplicity) derived from high-frequency (**A**-**B**) and low-frequency (**C**-**D**) ECoG features. Single-trial trajectories are shown for subjects S5 (78 trials; panel **A**,**C**) and S6 (73 trials; panel **B**,**D**). Inset boxes in each panel show the trajectories resulting from PCA performed directly on the ECoG features (without LFADS). The color code at bottom defines the portion of each trial corresponding to each behavioral mode.

In some cases, PCA-only resulted in a rough grouping of behavioral modes (pre-movement, movement, and force) into neural state space (ex. Figure 4A). However, the individual PCA-only trial trajectories remained highly variable, unlike the highly repeatable LFADS-PC trajectories. In other cases, PCA-only did not allow us to resolve a low-dimensional state space representation with identifiable groupings at all (ex. Figure 4D). Contrasting the LFADS-PC plots with the PCA-only plots (i.e., comparing each panel of Figure 4 with its inset) illustrates the benefit of LFADS on this dataset. We quantify this difference in Table 2, which shows the number of components required to account for 90% of the variance in the data, with and without LFADS.

**Table 2.**
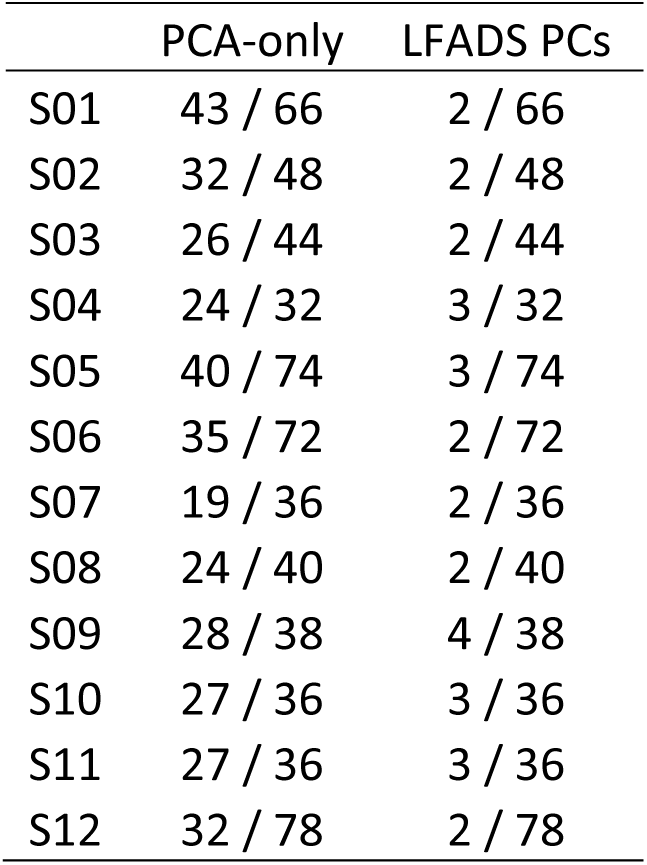
Number of principal components (PCs) required to account for 90% of the variance in the ECoG features (PCA-only) or the latent factors (LFADS PCs). Note that the number of features (factors) was equal to twice the number of ECoG electrodes selected for the analysis (those in M1/PM areas).

|  | PCA-only | LFADS PCs |
| --- | --- | --- |
| S01 | 43 / 66 | 2 / 66 |
| S02 | 32 / 48 | 2 / 48 |
| S03 | 26 / 44 | 2 / 44 |
| S04 | 24 / 32 | 3 / 32 |
| S05 | 40 / 74 | 3 / 74 |
| S06 | 35 / 72 | 2 / 72 |
| S07 | 19 / 36 | 2 / 36 |
| S08 | 24 / 40 | 2 / 40 |
| S09 | 28 / 38 | 4 / 38 |
| S10 | 27 / 36 | 3 / 36 |
| S11 | 27 / 36 | 3 / 36 |
| S12 | 32 / 78 | 2 / 78 |

Taken together, the results of Figure 4 and Table 2 illustrate the effectiveness of using LFADS to uncover low-dimensional representations of the neural state space during the kinematic-kinetic behavior. Examining the latent factors also provided strong additional evidence that the pre-movement, movement, and force behavioral modes were represented distinctly in the underlying ECoG signals.

### A neural vector angle summarizes temporal changes across the feature space

Visualizing the low-dimensional state space by LFADS-PCs reinforced the idea that pre-movement, movement, and force motor control modes are represented by distinct neural states. However, those methods did not allow us to generalize across subjects. Therefore, we used a second metric for summarizing the modulations of feature space across trials and subjects: the NVA. The NVA θ(t) is the angle at time t between a neural vector **m**(t) and its reference direction, **m**^ref^ (see Methods). Here, the high-dimensional vector **m**(t) was comprised of M1/PM ECoG spectral features. The reference vector **m**^ref^ was calculated during a window prior to the moment of peak force in each trial. Therefore θ(t) measures the dissimilarity between the ECoG features at each moment with their values during peak force generation.

To maximize the signal-to-noise ratio of θ(t), the elements of **m**(t) were selected using a cluster analysis (see Methods). In most cases, this approach resulted in (1) a cluster of well-modulated low-frequency features (ex. Figure 5A), (2) a cluster of well-modulated high-frequency features (ex. Figure 5B), and (3) a cluster of poorly modulated features (not shown). We computed θ(t) separately for clusters (1) and (2) in each subject (Figure 5C,D). The NVA recasts feature modulations for each trial into a common unit (angular difference in degrees). Therefore, we were able to combine NVA results across all trials in all subjects, yielding a compact study-wide representation of the cortical response to the movement-force transition (Figure 5E,F).

**Figure 5.**
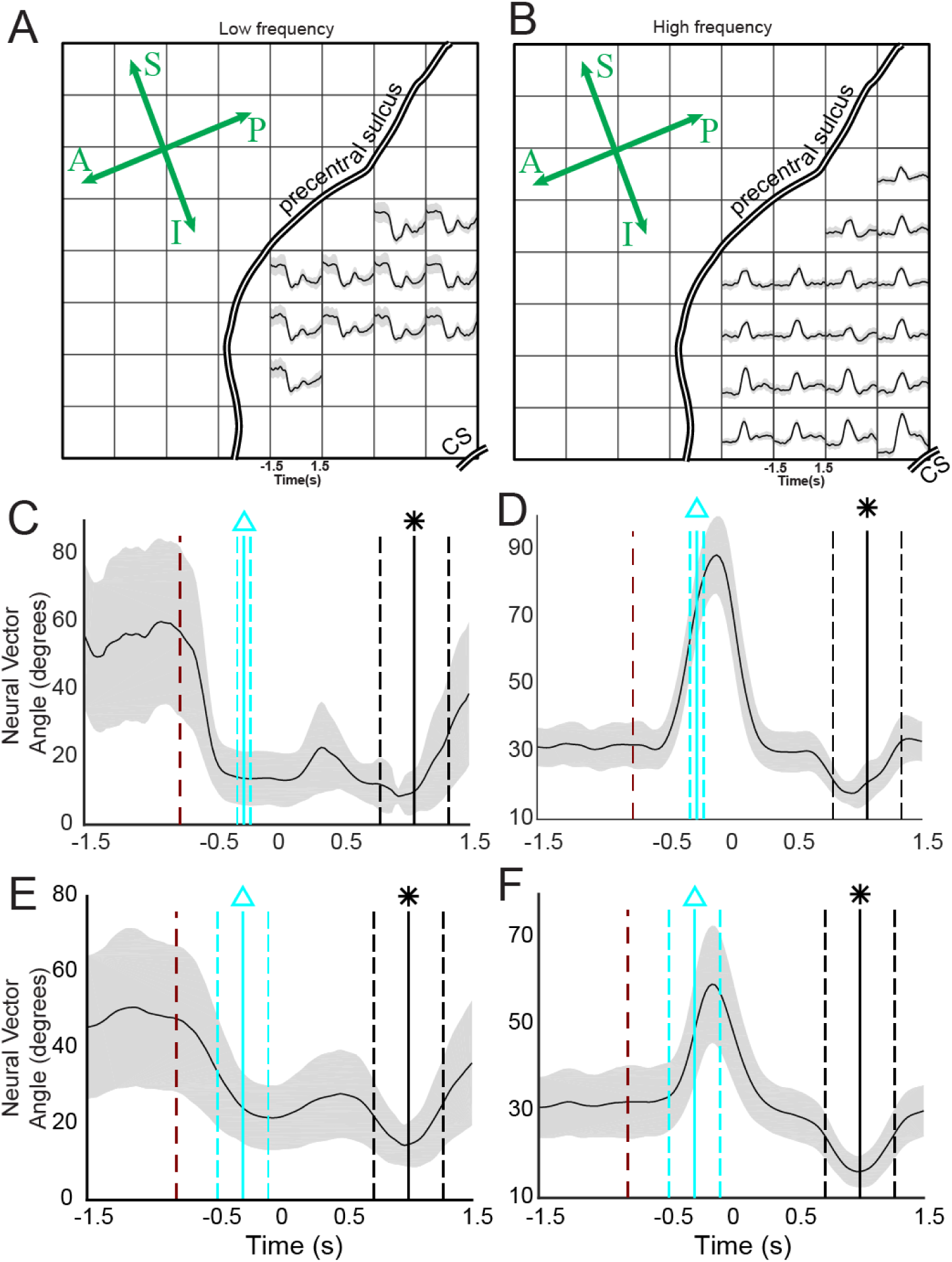
The neural vector angle (NVA) summarizes the cortical state change associated with the behavioral mode change from movement to force. (**A**,**B**) Electrodes selected for S5, using k-means clustering. CS; central sulcus. Anterior-posterior and superior-inferior are indicated on the rosette; compare to Figure 1A. (**A**) and (**B**) represent two of the three resulting clusters; the unsupervised cluster analysis natively divided the responses into low frequency and high frequency responses. (**C**) The NVA, θ(t) for the low frequency features selected in (**A)**. The dark red dashed line shows the average time of target appearance, relative to force onset (time=0). The vertical cyan lines show the mean (solid line) and standard deviation (dashed lines) of movement onset, relative to force onset. The vertical black lines show the time of maximum force for each trial (equivalent to the reference period **m**^ref^). (**D**) The NVA for the high frequency features shown in (**B**). (**E**,**F**) NVAs calculated across all trials, all subjects in the study. Labeling conventions are the same as in (**C**,**D**).

Across subjects, average low-frequency NVAs began to decrease immediately after the presentation of the force target (Figure 5E, red line), and reached their minimum value approximately at the start of flexion (Figure 5E, cyan line). Accordingly, low-frequency NVA during movement was significantly lower than NVA during the pre-movement period (p<10^-9^; Kruskal-Wallis test, Tukey HSD post-hoc for all statistical comparisons in this section). By contrast, there was no significant difference between the movement period and force (t=0 to t=0.75) in the low-frequency NVAs (p=0.32). High-frequency NVAs did not deviate from their pre-movement values at target presentation (Figure 5F), instead changing just prior to the start of movement (Figure 5F, cyan line). During movement, high-frequency NVAs were significantly higher than pre-movement NVA (p<10^-9^), peaking just before the onset of force (Figure 5F, approximately t= −130 ms relative to force onset). During the force behavioral mode, high-frequency NVA were overall lower than either movement (p<10^-9^) or pre-movement (p<10^-6^) periods.

Overall, the NVA results indicate that separate cortical states are responsible for pre-movement, movement, and force behavioral modes. In addition, we found evidence for a possible distinction in roles, or kinds of information encoded by low- versus high-frequency ECoG features. This was illustrated by the fact that low-frequency NVAs did not differentiate force and movement, while high frequency NVAs did differentiate those two behavioral modes. Earlier, we found evidence for encoding multiple types of information on an example electrode (Figure 2B,E), which modulated its spectral intensity differently in low- versus high-frequency spectral domains. Here, the NVA results provide evidence that this may be a general feature of PM/M1 cortical areas.

### ECoG features enabled accurate classification of behavioral modes

The above evidence indicates that during grasp, the behavioral modes of finger movement and force are represented by distinct neural states in the motor and premotor cortices. This has potential applications for brain-machine interface (BMI) design. For example, in response to changing functional goals (e.g., changing from movement control to force control when picking up an object), a BMI could switch control strategies. To estimate the accuracy such control might achieve, we tested whether the subjects’ behavioral mode could be decoded from cortical activity. We used the low- and high-frequency ECoG spectral features to classify each time bin as one of three behavioral modes: pre-movement, movement, or force execution. In parallel with the ECoG feature-based classification, we also classified behavioral mode using the LFADS-denoised features as inputs. We used two widely available classifiers: support vector machines (SVM) and boosted aggregate (bagged) decision trees. For each subject, we also calculated a chance decoding value (see Methods). We report classification accuracy for the two types of classifiers separately, evaluating both the features and the LFADS-denoised features. The three behavioral modes were strongly differentiable in all subjects, with high accuracy (Figure 6). Overall, the tree-based classifier outperformed SVM, and LFADS-denoised features were decoded more accurately than the features without denoising (p=1.9^-7^, Kruskal-Wallis test). For the tree-based classifier of LFADS-denoised features, median decoding accuracies for the subjects ranged from 87%±2% to 94%±1%, with an overall median value of 90%±6%, indicating that these three classes were highly separable. Statistically, the decoding accuracy for all subjects was significantly higher than chance. Thus, these behavioral modes have highly separable cortical representations.

**Figure 6.**
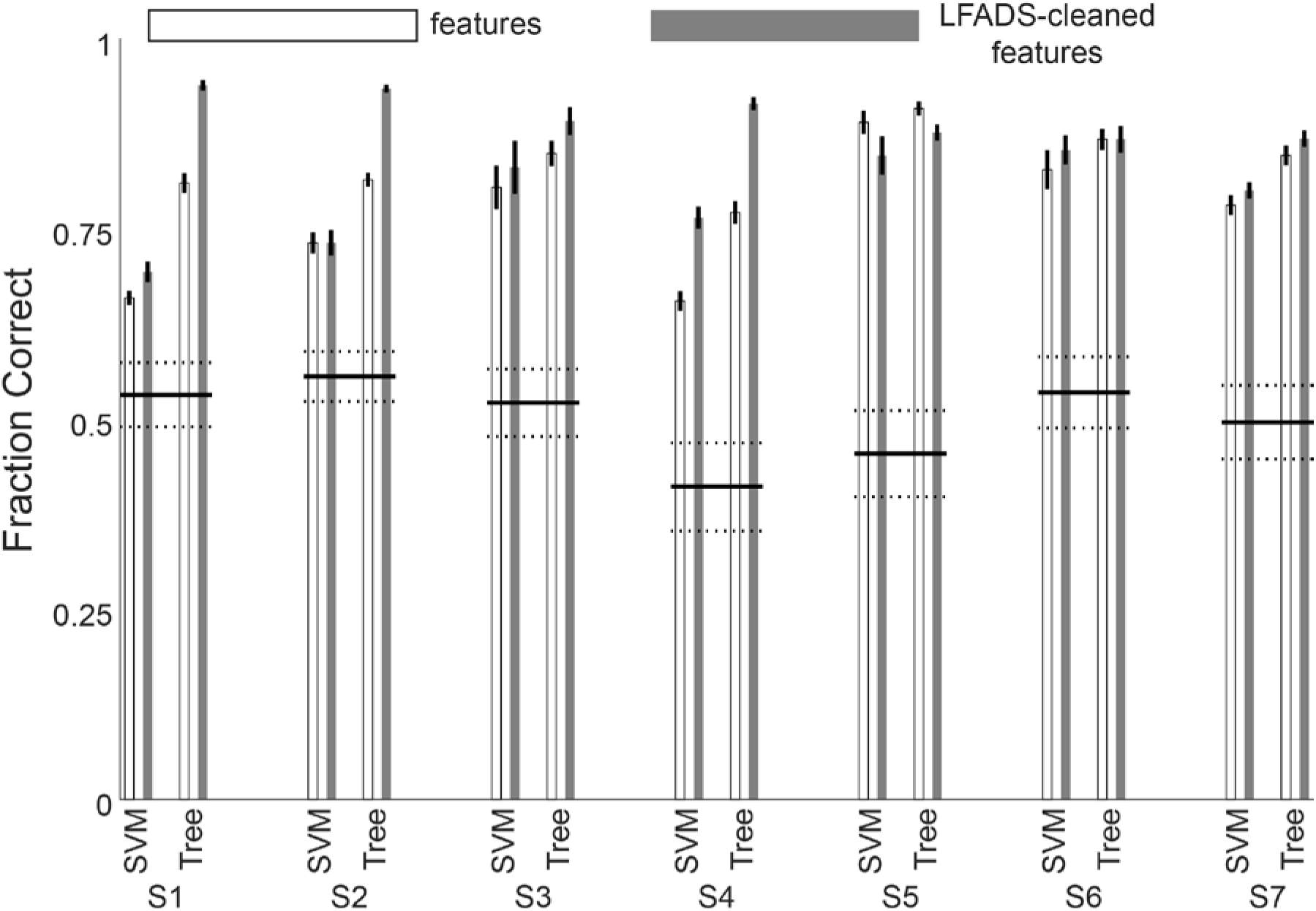
Decoding behavioral mode from ECoG features before and after LFADS denoising. The median classification accuracy was greater than chance for all subjects. SVM; support vector machines. Tree; boosted aggregate decision tree classifier.

## Discussion

Manipulating objects dexterously requires controlling both grasp kinematics and isometric force. Even simple activities like turning a doorknob, shaking hands, and lifting a cup of liquid could not be accomplished safely and quickly without both kinds of control. More than two decades ago, investigators began to appreciate that the cortex may handle these two vital aspects of motor behavior separately (Flanagan et al., 1999). Here, we found distinct and quantifiable differences in how the motor and premotor cortices represented behavioral mode, i.e. pre-movement, flexion movement and isometric force. Notably, low-frequency ECoG features seemed to modulate their activity with movement onset, while high-frequency ECoG features often modulated with force onset. Feature modulations were time-locked to behaviorally relevant events, and could be detected on a-single trial basis (Figure 2). The ensemble ECoG modulations constituted a neural state change, accompanying the changes in behavioral mode (from pre-movement to movement, or from movement to force). We were able to model this change using a dynamical systems approach (LFADS), and decode the subjects’ behavioral modes with high accuracy. Understanding neural state changes like these in the context of a functional grasp task has implications for the design of dexterous grasp brain-machine interfaces.

As in previous work, we decoded the continuous time course of the behavioral variables (movement and force). Generally, we achieved highly accurate decoding of both force and movement, comparing favorably with prior studies decoding finger movement kinematics (Acharya et al., 2010; Nakanishi et al., 2014; Xie et al., 2018) and isometric force (Pistohl et al., 2013; Chen et al., 2014; Flint et al., 2014). Most importantly, there was no significant difference in our ability to decode force and movement across subjects. This implies that the differences in cortical representations of force and movement were not simply expressions of a superior decoding of one or the other.

Spatially, human cortical encoding of finger movement takes place over a widespread area (Schieber, 2002), including complex and overlapping representations of individual finger movements (Dechent and Frahm, 2003). ECoG recordings enabled us to examine cortical activity on these relatively large spatial scales. We found that the maps of decoding performance altered significantly across movement and force representations (across-mode) in 6 of 7 subjects. We controlled for changes due to time or behavioral variability (within-mode), by comparing the between-mode maps to the within-mode maps. One potential explanation for the spatial map differences could be that the activating regions of the maps are simply shrinking during isometric force. Such an explanation is consistent with evidence pointing to less cortical modulation with isometric force than with movement (Hendrix et al., 2009). However, in this case we found that the peaks of the decoding maps changed location (Table 1), indicating that the maps shifted rather than merely growing or shrinking. These spatial decoding results are relevant to the design of brain-machine interfaces (BMIs), since any BMI that restores grasp should ideally execute both movement and force functions. There is evidence that representations of hand movements are preserved following amputation (Bruurmijn et al., 2017), though it remains to be shown whether the movement-force functional map change will remain in an individual with paralysis. Downey et al. (2017) found that applying a scaling factor to neuronal spike rates facilitated the ability of human BMI users to grasp objects with a prosthetic hand. The utility of such a scaling factor may be a reflection of the functional somatotopy of the cortex, though the current results suggest that amplitude scaling would not necessarily be the ideal method of accounting for the difference in movement and force representations.

Increasingly, spiking activity in small areas of motor cortex have been modeled as dynamical systems in an effort to parsimoniously describe and understand their network-level activity. In this study, we used PCA to visualize the low-dimensional neural state space LFADS uncovered for each subject. The LFADS-PCs were tightly grouped over trials and occupied distinct regions of state space during the pre-movement, movement, and force behavioral modes (Figure 4B,D). Both low-frequency and high-frequency LFADS-PCs were clearly separated in different behavioral modes. Some previous examples of modeling cortical dynamics using latent factors have analyzed single motor control modes. For example, Vaidya et al. (2015) modeled both reach- and grasp-related neural ensembles as linear dynamical systems to study learning. Also, Gallego et al. (2018) also showed that there were some differences in local M1 neuronal ensemble activity between kinematic and kinetic cursor control tasks. Our results show that dynamical systems modeling can elucidate the latent factors underlying a widespread cortical network in addition to local circuit networks. It was not surprising that latent factor state space trajectories evolved with time during each trial; indeed, this is a fundamental underlying assumption of the dynamical systems model. The significance of the LFADS-derived trajectories was their smooth, repeatable paths through distinct regions of state space during behavioral mode transitions. Compared with PCA-only state space trajectories, LFADS factors clustered more tightly and evolved much more repeatably in pre-movement, movement, and force behavioral modes.

We used the NVA to summarize spectro-temporal changes across electrodes and subjects. We found that NVAs from low-frequency features changed with the start of movement. NVAs from high-frequency features changed between movement and force control modes. The average duration of high-frequency neural vector changes (about 300 ms; Figure 5F) was substantially shorter than the average duration of the force-matching part of the behavioral task (about 1 s). This profile of activation (phasic rise in high gamma modulation near the onset of behavior) has been shown during isotonic movement as well (Flint et al., 2017). It appears that the onset of force control, or perhaps the transition from movement to force, is especially meaningful to the cortex when encoding grasp.

Our results, particularly the NVA analysis, support and extend the findings of Venkadesan and Valero-Cuevas (2008), who inferred from muscle activity that the human motor system uses two separate control strategies for movement and isometric force. Importantly, this muscle activity changed about 100 ms before force onset, ruling out the conclusion that changes in EMG patterns are purely the result of the mechanical constraints of the behavior. In the current three-behavioral-mode paradigm (pre-movement, movement, and force), we found that low-frequency NVA clearly differentiated movement from pre-movement, but failed to differentiate movement from force. High-frequency NVA allowed us to differentiate all three behavioral modes. Together, these results provide direct evidence of separable cortical representations for movement and force. Moreover, the change in high gamma activity patterns (reflected by the NVA) occurred around 130 ms prior to force onset. This time course of changing cortical activity is consistent with the earlier EMG results, and with the concept that control strategies for movement and force are encoded in the motor and premotor cortices, rather than subcortical systems. This argues against the hypothesis that differences in cortical activity during movement-force are due mainly to somatosensory feedback changes in the two states.

Our decoding of the subjects’ time-varying behavioral mode has ramifications for BMI design, as demonstrated by Suminski et al. (2013). Suminski et al. addressed a longstanding limitation of BMIs: decoders trained on a given set of motor activities do not predict accurately outside those activities. Hierarchical BMIs, which include multiple decoders operating in parallel with a switching mechanism, are likely to outperform single-decoder BMIs. Given the differences we observed in movement and force representation, it seems unlikely that a decoder trained only on grasping movement data will provide optimal control of a BMI for grasping and manipulating objects, either with a prosthetic hand or functional electrical stimulation of paralyzed fingers. Our results suggest decoding the behavioral mode from cortical activity is feasible and could increase the functionality of BMIs during object grasp. The improvements in behavioral mode decoding by using latent factors indicates that viewing the cortical motor control circuits as a dynamical system can facilitate the task of identifying cortical correlates of multiple behavioral modes.

## Acknowledgements

We would like to thank our research subjects for their participation, and Mukta Vaidya for helpful comments on the manuscript. This work was supported by the following sources: Craig H Neilsen Foundation fellowship (RDF); Emory College Computational Neuroscience training grant (KL); Burroughs Wellcome Fund Collaborative Research Travel Grant (CP); National Science Foundation NCS 1835364 (CP); Emory Neuromodulation Technology Innovation Center (CP); Doris Duke Charitable Foundation Clinical Scientist Development Award (MWS); Northwestern Memorial Foundation Dixon Translational Research Grant Program (supported in part by NIH Grant UL1RR025741) (MWS); National Institutes of Health R01NS094748 (MWS).

## References

1. Acharya S, Fifer MS, Benz HL, Crone NE, Thakor NV (2010) Electrocorticographic amplitude predicts finger positions during slow grasping motions of the hand. J Neural Eng 7:046002.

2. Ajiboye AB, Willett FR, Young DR, Memberg WD, Murphy BA, Miller JP, Walter BL, Sweet JA, Hoyen HA, Keith MW, Peckham PH, Simeral JD, Donoghue JP, Hochberg LR, Kirsch RF (2017) Restoration of reaching and grasping movements through brain-controlled muscle stimulation in a person with tetraplegia: a proof-of-concept demonstration. Lancet 389:1821–1830.

3. Blabe CH, Gilja V, Chestek CA, Shenoy KV, Anderson KD, Henderson JM (2015) Assessment of brain-machine interfaces from the perspective of people with paralysis. J Neural Eng 12:043002.

4. Bouton CE, Shaikhouni A, Annetta NV, Bockbrader MA, Friedenberg DA, Nielson DM, Sharma G, Sederberg PB, Glenn BC, Mysiw WJ, Morgan AG, Deogaonkar M, Rezai AR (2016) Restoring cortical control of functional movement in a human with quadriplegia. Nature 533:247–250.

5. Bruurmijn M, Pereboom IPL, Vansteensel MJ, Raemaekers MAH, Ramsey NF (2017) Preservation of hand movement representation in the sensorimotor areas of amputees. Brain 140:3166–3178.

6. Chen C, Shin D, Watanabe H, Nakanishi Y, Kambara H, Yoshimura N, Nambu A, Isa T, Nishimura Y, Koike Y (2014) Decoding grasp force profile from electrocorticography signals in non-human primate sensorimotor cortex. Neurosci Res 83:1–7.

7. Chib VS, Krutky MA, Lynch KM, Mussa-Ivaldi FA (2009) The separate neural control of hand movements and contact forces. J Neurosci 29:3939–3947.

8. Churchland MM, Cunningham JP, Kaufman MT, Foster JD, Nuyujukian P, Ryu SI, Shenoy KV (2012) Neural population dynamics during reaching. Nature 487:51–56.

9. Collard MJ, Fifer MS, Benz HL, McMullen DP, Wang Y, Milsap GW, Korzeniewska A, Crone NE (2016) Cortical subnetwork dynamics during human language tasks. Neuroimage 135:261–272.

10. Danion F, Diamond JS, Flanagan JR (2013) Separate contributions of kinematic and kinetic errors to trajectory and grip force adaptation when transporting novel hand-held loads. J Neurosci 33:2229–2236.

11. Dechent P, Frahm J (2003) Functional somatotopy of finger representations in human primary motor cortex. Hum Brain Mapp 18:272–283.

12. Downey JE, Brane L, Gaunt RA, Tyler-Kabara EC, Boninger ML, Collinger JL (2017) Motor cortical activity changes during neuroprosthetic-controlled object interaction. Sci Rep 7:16947.

13. Evarts EV (1968) Relation of pyramidal tract activity to force exerted during voluntary movement. J Neurophysiol 31:14–27.

14. Fagg AH, Ojakangas GW, Miller LE, Hatsopoulos NG (2009) Kinetic trajectory decoding using motor cortical ensembles. IEEE Trans Neural Syst Rehabil Eng 17:487–496.

15. Flanagan JR, Nakano E, Imamizu H, Osu R, Yoshioka T, Kawato M (1999) Composition and decomposition of internal models in motor learning under altered kinematic and dynamic environments. J Neurosci 19:RC34.

16. Flint RD, Rosenow JM, Tate MC, Slutzky MW (2017) Continuous decoding of human grasp kinematics using epidural and subdural signals. J Neural Eng 14:016005.

17. Flint RD, Wang PT, Wright ZA, King CE, Krucoff MO, Schuele SU, Rosenow JM, Hsu FP, Liu CY, Lin JJ, Sazgar M, Millett DE, Shaw SJ, Nenadic Z, Do AH, Slutzky MW (2014) Extracting kinetic information from human motor cortical signals. Neuroimage 101:695–703.

18. Gallego JA, Perich MG, Naufel SN, Ethier C, Solla SA, Miller LE (2018) Cortical population activity within a preserved neural manifold underlies multiple motor behaviors. Nat Commun 9:4233.

19. Hendrix CM, Mason CR, Ebner TJ (2009) Signaling of grasp dimension and grasp force in dorsal premotor cortex and primary motor cortex neurons during reach to grasp in the monkey. J Neurophysiol 102:132–145.

20. Hochberg LR, Bacher D, Jarosiewicz B, Masse NY, Simeral JD, Vogel J, Haddadin S, Liu J, Cash SS, van der Smagt P, Donoghue JP (2012) Reach and grasp by people with tetraplegia using a neurally controlled robotic arm. Nature 485:372–375.

21. Hotson G, McMullen DP, Fifer MS, Johannes MS, Katyal KD, Para MP, Armiger R, Anderson WS, Thakor NV, Wester BA, Crone NE (2016) Individual finger control of a modular prosthetic limb using high-density electrocorticography in a human subject. J Neural Eng 13:026017.

22. Hunter IW, Korenberg MJ (1986) The identification of nonlinear biological systems: Wiener and Hammerstein cascade models. Biol Cybern 55:135–144.

23. Moran DW, Schwartz AB (1999) Motor cortical representation of speed and direction during reaching. J Neurophysiol 82:2676–2692.

24. Nakanishi Y, Yanagisawa T, Shin D, Chen C, Kambara H, Yoshimura N, Fukuma R, Kishima H, Hirata M, Koike Y (2014) Decoding fingertip trajectory from electrocorticographic signals in humans. Neurosci Res 85:20–27.

25. Pandarinath C, O’Shea DJ, Collins J, Jozefowicz R, Stavisky SD, Kao JC, Trautmann EM, Kaufman MT, Ryu SI, Hochberg LR, Henderson JM, Shenoy KV, Abbott LF, Sussillo D (2018) Inferring single-trial neural population dynamics using sequential auto-encoders. Nature methods 15:805–815.

26. Pfurtscheller G, Muller GR, Pfurtscheller J, Gerner HJ, Rupp R (2003) ’Thought’-- control of functional electrical stimulation to restore hand grasp in a patient with tetraplegia. Neurosci Lett 351:33–36.

27. Pistohl T, Schmidt TS, Ball T, Schulze-Bonhage A, Aertsen A, Mehring C (2013) Grasp Detection from Human ECoG during Natural Reach-to-Grasp Movements. PLoS One 8:e54658.

28. Schalk G, McFarland DJ, Hinterberger T, Birbaumer N, Wolpaw JR (2004) BCI2000: a general-purpose brain-computer interface (BCI) system. IEEE Trans Biomed Eng 51:1034–1043.

29. Schieber MH (2002) Motor cortex and the distributed anatomy of finger movements. Adv Exp Med Biol 508:411–416.

30. Suminski AJ, Tkach DC, Fagg AH, Hatsopoulos NG (2010) Incorporating feedback from multiple sensory modalities enhances brain-machine interface control. J Neurosci 30:16777–16787.

31. Suminski AJ, Fagg AH, Willett FR, Bodenhamer M, Hatsopoulos NG (2013) Online adaptive decoding of intended movements with a hybrid kinetic and kinematic brain machine interface. Conf Proc IEEE Eng Med Biol Soc 2013:1583–1586.

32. Sussillo D, Jozefowicz R, Abbott L, Pandarinath C (2016) Lfads-latent factor analysis via dynamical systems. arXiv preprint arXiv:160806315.

33. Vaidya M, Kording K, Saleh M, Takahashi K, Hatsopoulos NG (2015) Neural coordination during reach-to-grasp. J Neurophysiol 114:1827–1836.

34. Venkadesan M, Valero-Cuevas FJ (2008) Neural control of motion-to-force transitions with the fingertip. J Neurosci 28:1366–1373.

35. Wang PT, King CE, McCrimmon CM, Lin JJ, Sazgar M, Hsu FP, Shaw SJ, Millet DE, Chui LA, Liu CY, Do AH, Nenadic Z (2016) Comparison of decoding resolution of standard and high-density electrocorticogram electrodes. J Neural Eng 13:026016.

36. Wodlinger B, Downey JE, Tyler-Kabara EC, Schwartz AB, Boninger ML, Collinger JL (2014) Ten-dimensional anthropomorphic arm control in a human brain-machine interface: difficulties, solutions, and limitations. J Neural Eng 12:016011.

37. Xie Z, Schwartz O, Prasad A (2018) Decoding of finger trajectory from ECoG using deep learning. J Neural Eng 15:036009.

38. Yanagisawa T, Hirata M, Saitoh Y, Kishima H, Matsushita K, Goto T, Fukuma R, Yokoi H, Kamitani Y, Yoshimine T (2012) Electrocorticographic control of a prosthetic arm in paralyzed patients. Annals of neurology 71:353–361.

